# Hormonal regulation of social ascent and temporal patterns of behavior in an African cichlid

**DOI:** 10.1101/414045

**Authors:** Beau A. Alward, Austin T. Hilliard, Ryan A. York, Russell D. Fernald

## Abstract

For many species, social rank determines which individuals perform certain social behaviors and when. Higher ranking or dominant (DOM) individuals maintain status through aggressive interactions and perform courtship behaviors while non-dominant (ND) individuals do not. In some species ND individuals ascend (ASC) in social rank when the opportunity arises. Many important questions related to the mechanistic basis of social ascent remain to be answered. We probed whether androgen signaling regulates social ascent in male *Astatotilapia burtoni*, an African cichlid whose social hierarchy can be readily controlled in the laboratory. As expected, androgen receptor (AR) antagonism abolished reproductive behavior during social ascent. However, we discovered multiple AR-dependent—and AR-independent—temporal behavioral patterns that typify social ascent and dominance. AR antagonism in ASC males reduced the speed of behavioral performance compared to DOM males. Socially ascending males, independent of AR activation, were more likely than DOM males to follow aggressive displays with another aggressive display. Further analyses revealed differences in the sequencing of aggressive and courtship behaviors, wherein DOM males were more likely than ASC males to follow male-directed aggression with courtship displays. Strikingly, this difference was driven mostly by ASC males taking longer to transition from aggression to courtship, suggesting ASC males can perform certain DOM-typical temporal behavioral patterns. Our results indicate androgen signaling drives social ascent, but hormonal signaling and social experience shape the full suite of DOM-typical behavioral patterns.

## Introduction

Social animals often organize into hierarchies, where higher ranking members have access to key resources such as territory, food, and mates (Sapolsky, 2005; Wilson, 1975). Such hierarchies are established through agonistic interactions between conspecifics. Higher ranking individuals perform a variety of courtship behaviors that are part of the species-typical suite of acts that culminate in copulation and reproduction (Adkins-Regan, 2009).

Studies in primates, birds, and fish have discovered key social and environmental factors that regulate social status (Fernald, 2012; Sapolsky, 2005). The African cichlid fish *Astatotilapia burtoni* has been used as a particularly popular model for the study of social behavior given the unique opportunities if offers for mechanistic analysis of status. In their native habitat, Lake Tanganyika, *A. burtoni* males exist as either dominant (DOM) or non-dominant (ND; Figure 1A). ND males survey the social environment waiting for an opportunity to rise in social rank, a process that has been called social ascent, wherein males exhibit an increase in aggressive and reproductive behaviors typical of DOM males (Figure 1A-B; Burmeister et al., 2005; Maruska, 2015; Maruska et al., 2013; Maruska and Fernald, 2013, 2010). DOM males possess large gonads and high circulating levels of testosterone and its androgenic and estrogenic metabolites, while ND males have small gonads and low levels of circulating testosterone. Ascending (ASC) males experience a large increase in circulating testosterone to DOM-typical levels, even though their gonads remain small. Aggression in *A. burtoni* appears to be mediated by the actions of estrogenic metabolites of testosterone (Hufman et al., 2013; O’Connell and Hofmann, 2012a). However, the role of androgen signaling in mediating social behavior in *A. burtoni* is still unclear, with contradictory results in the literature. Blocking androgen receptors (AR) with the potent antagonist cyproterone acetate (CA) can indeed reduce specific courtship behaviors in stable DOM males (O’Connell and Hofmann, 2012a), yet injection of dihydrotestosterone, an androgen, fails to enhance courtship in ND males. Thus, while estrogen signaling regulates aggression regardless of social status, the role of androgen signaling is less clear. For example, how does an ND male, when given the opportunity, exhibit an increase in an AR-dependent behavior, courtship?

**Figure 1.**
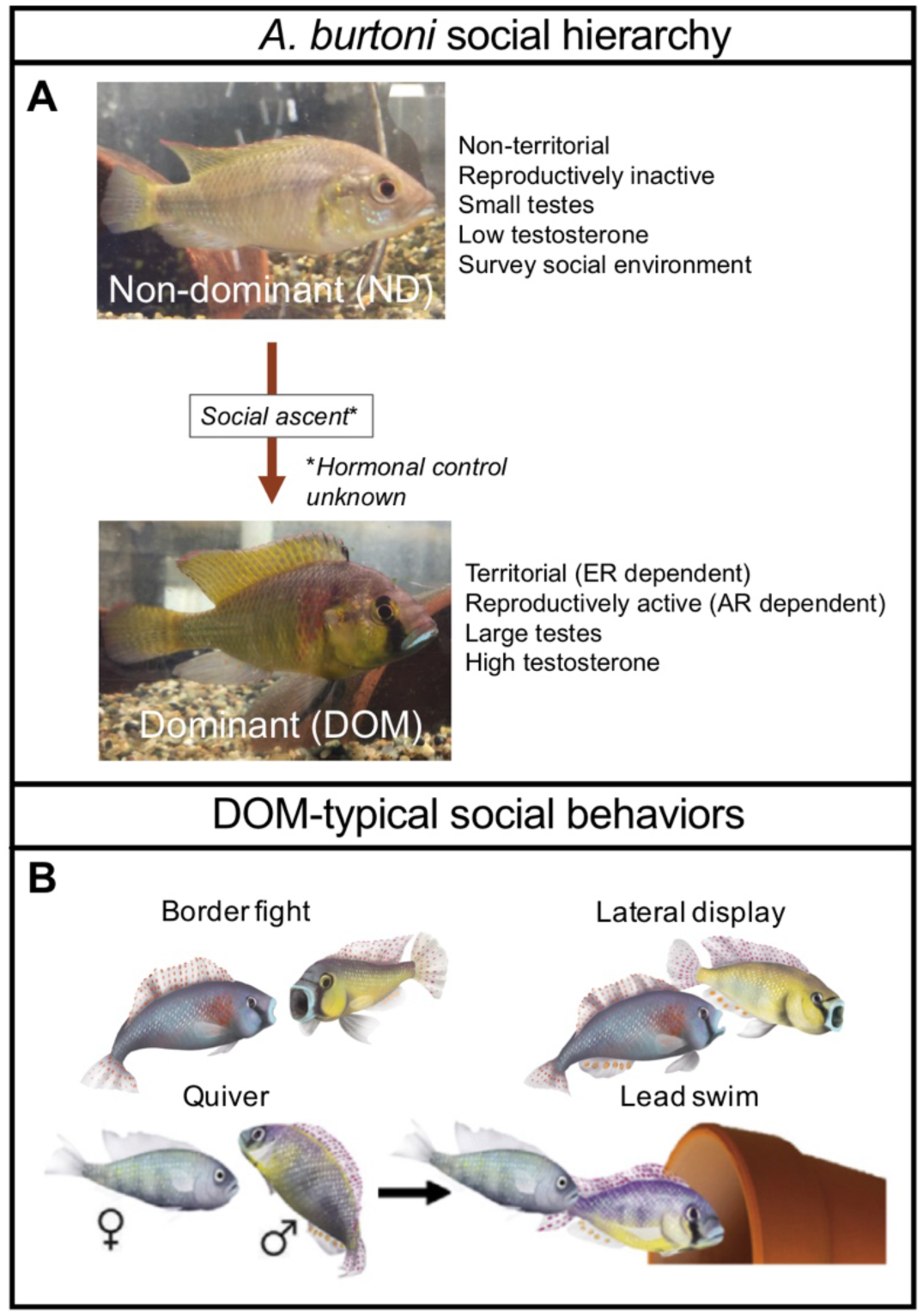
*A. burtoni* males exist in a social hierarchy, in which (A) ND males survey the social environment and attempt to rise to DOM male status when the opportunity arises (Fernald, 2017); (B) DOM males exhibit a complex suite of aggressive (e.g., border fight and lateral display) and reproductive (e.g., quiver and lead swim) behaviors that are readily quantifiable in naturalistic laboratory settings. ER=Estrogen receptor; AR=Androgen receptor.

Seasonally-breeding songbirds, a model system in the hormonal control of behavior, may provide clues (Alward et al., 2018). Songbirds undergo a transformation in courtship behavior as they transition from non-breeding to breeding conditions, a process that could be viewed as similar to social ascent. For instance, when in non-breeding conditions (e.g., during the fall and winter), male canaries (*Serinus canaria)* and white crown sparrows (*Zonotrichea leucophrys*) possess small testes, low testosterone, and sing low levels of courtship song (Reviewed in Alward et al., 2018 and Schlinger and Brenowitz, 2009). However, when in breeding conditions (e.g., during the spring), males experience a dramatic increase in testosterone levels and courtship song, which is followed several days later by an increase in testes size. Laboratory studies have found the increase in courtship song is AR- and estrogen receptor (ER)-dependent (Reviewed in Ball et al., 2002 and Schlinger and Brenowitz, 2009). Moreover, global and localized manipulations using pharmacological techniques have found AR activation enhances courtship song (Sartor et al., 2005), but also controls multiple temporal aspects of song, such as the tempo of specific vocalizations and their sequential arrangement (Alward et al., 2017).

Do ascending *A. burtoni* males undergo similar AR-dependent shifts in the activation and temporal sequencing of specific behaviors? Here, we address this question via an investigation of social ascent across behavioral and temporal scales. To do so we augment standard measures of DOM-typical aggressive and courtship behaviors (Fernald and Maruska, 2012; Maruska and Fernald, 2013) with novel assays of behavioral sequence and interval variation across time scales relevant to ascent. We initially found blockade of AR abolished the rise in reproductive behavior normally seen in ASC fish, as expected given previous observations. However, upon factoring in temporal information, it was revealed that AR activation is also required for enhancing the speed of behavioral performance during ascent. Further investigation into these temporal patterns of behavior revealed that socially ascending fish—regardless of AR activation—significantly diverged from stable DOM males in the sequelae of behavior at multiple time-scales. For instance, socially ascending males were more likely than stable DOM males to perform aggressive displays twice in a row, while DOM males were significantly more likely than socially ascending males to follow aggressive behavior with courtship behavior. Strikingly, several of these differences disappeared when we used a longer interval for defining bouts of behavior, suggesting ASC males can perform DOM-like behavior sequencing patterns but take longer to do so. Our findings expand our knowledge of the hormonal control of social status and, to best of our knowledge, identify for the first time that temporal patterns of behavior over multiple time-scales are key factors in demarcating social status. Furthermore, our data suggest social behavior in *A. burtoni* is highly dissociable, with motivation and temporal aspects of behavior likely being mediated by independent mechanisms.

## Materials and Methods

### Animals

Adult *A. burtoni* were derived from wild-caught stock from Lake Tanganyika, Africa, and laboratory-bred in aquaria under environmental conditions that mimic their natural equatorial habitat (28 °C; pH 8.0; 12:12 h light/dark cycle with full spectrum illumination; constant aeration). Aquaria contained gravel-covered bottoms with terra cotta pots cut in half to serve as shelters and spawning territories. Fish were fed cichlid pellets and flakes (AquaDine, Healdsburg, CA, USA) each morning. All experimental procedures were approved by the Stanford Administrative Panel for Laboratory Animal Care.

### Social manipulation and CA injection

#### Rationale

*A. burtoni*, like many other teleosts, have two ARs (Harbott et al., 2007). While ND males have lower levels of circulating androgens compared to DOM males, they are not undetectable (Parikh et al., 2006). Androgens could promote social ascent in two ways: low levels of androgens in ND males promote ascent once the opportunity arises and/or androgens act rapidly on the day of ascent promoting a rise in social status. Thus, in our injection procedures we aimed to address these two issues. To block ARs, we injected the AR antagonist cyproterone acetate (CA), a potent steroidal anti-androgen that blocks androgen binding to the ligand binding domain of ARs through competitive binding (Fourcade and McLeod, 2004). Previous work in *A. burtoni* using CA showed that maximal behavioral effects were observed two days after injection, but significant behavioral effects occurred on the day of and day after injection (O’Connell and Hofmann, 2012a). Thus, our injection procedure likely blocks ARs on the day of injection up until the day of ascent.

#### Experimental design

We used a previously described “social ascent” paradigm (Maruska et al., 2011; Maruska and Fernald, 2011, 2010) that is shown in Figure 2. Briefly, to establish socially-suppressed ND fish, males were placed into aquaria for 4–5 weeks with several larger dominant suppressor males, females, and ND males. At the end of the suppression period, subjects were moved into the central compartment of an experimental tank that contained one larger resident dominant male and three females. This central compartment was separated with transparent acrylic barriers from flanking compartments that contained two smaller males and three females of various reproductive states so that fish could interact visually but not physically with community animals. After an hour acclimation, we recorded fish from 1:30 pm to 2:00 pm (see Recording and scoring of behavior below). Then, the focal male was removed, injected intraperitoneally with CA (for a subset of ND males that would be allowed to socially ascend, hereon called ASC+CA; concentration of CA=0.83 µg/gbw) or vehicle (sesame oil; given to DOM, ASC, and ND fish) and immediately placed back into the central compartment of the experimental tank. The concentration of CA was determined based on previous work showing that injecting CA at this concentration results in significant behavioral effects two days after injection (O’Connell and Hofmann, 2012). Two days later—the day of ascent—the larger resident suppressor male was removed with a net 30 minutes before light onset using infrared night vision goggles (Model 26-1020; Bushnell, Overland Park, KS, USA), providing an opportunity for the ND male to ascend at light onset. Stable DOM and stable ND males were used as controls and compared to males ascending in social status. Stable ND males were suppressed in community tanks for 4–5 weeks and transferred to the same experimental tank as described above. On the day of ascent, however, removal of the suppressor DOM male was only simulated by dipping a net into the tank before light onset without removal of the dominant resident. Stable DOM were DOM males that maintained their status in community tanks for 4–5 weeks and were then placed in the experimental tank with females but no larger resident male, which maintains their dominance status. On the day of ascent, a net was dipped into the water before light onset to simulate resident male removal.

**Figure 2.**
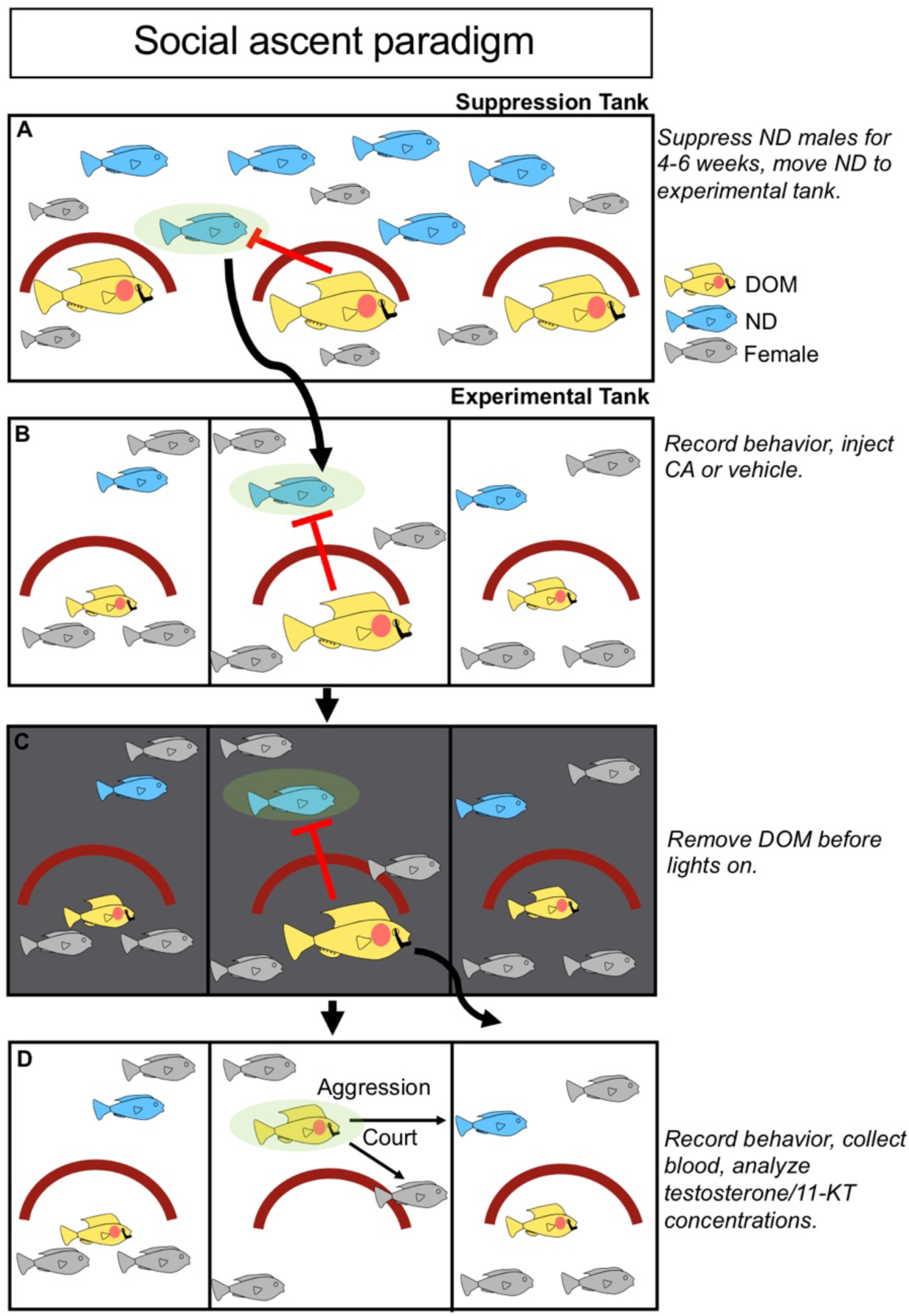
Schematic showing the social ascent paradigm combined with the injection of CA or vehicle. Males were placed into community suppression tanks for 4– 5 weeks that contained three large dominant males (i.e., suppressor males) and females. (B) Suppressed males were then transferred to a central compartment of an experimental tank for 2 days prior to social opportunity. The central compartment contained a larger resident dominant male and 3 females and was separated with transparent acrylic barriers on either side from community tanks containing females, subordinate males, and dominant males that were smaller in size than the subject fish. On the day of transfer to the experimental tank, fish were allowed to acclimate for 1 hour; then, behavior was recorded for 30 minutes and fish were injected with either vehicle (sesame oil) or CA, an AR antagonist. Fish were filmed again the following day. (C) Then, on the third day (i.e. the day of social ascent) the resident dominant male was removed from the central compartment 1 h before the lights turned on in the morning. (D) At light onset, behavior was filmed for five hours and fish were removed and blood was collected for analyses of concentrations of the androgens testosterone and 11-KT. Upside down red “U”s indicate the halved terra cotta pots that functioned as the males’ spawning sites. DOM=dominant; ND=non-dominant. KT=ketotestosterone.

### Recording and scoring of behavior

Behavior was quantified during essential 30-minute intervals: after an hour of acclimation on day 1, but immediately before the injection (“pre-injection”) from 1:30 pm to 2:00 pm; on day 2 from 1:30 to 2:00 pm (baseline (BL)); and on day 3 (day of ascent) from 9:00 am to 9:30 am and 1:30 to 2:00 pm. Focal fish were removed from the experimental compartment at 2:00 pm on the day of ascent, when tissue was harvested for physiological measurements and blood collection (see Morphological and steroid hormone analyses). Behavior was recorded using a digital video camera (Sony, HDR). Videos were scored by an observer without knowledge of the fish identification number. Multiple behaviors were quantified (Fernald and Hirata, 1977): subordinate behavior (flee); territorial or agonistic behaviors (lateral display and border fight); and reproductive behaviors (chase female, courtship display, lead swim, pot entry, and dig). Fleeing was defined as a rapid retreat swim from an approaching dominant male or female. Lateral displays were classified as presentations of the side of the body to another fish with erect fins, flared opercula, and body quivering. Border fights were interactions with neighboring dominant males across the acrylic barrier typified by head-on lunges and rams against the barrier with an open mouth. Chase female was defined as a rapid swim directed towards a female and was grouped with reproductive behaviors here because they were only directed towards females within the same compartment, and chases are a normal component of the courtship repertoire. Courtship displays were defined as rapid quivers by the male with presentation of the anal fin egg spots, and lead swim was defined as swimming towards the shelter accompanied by back-and-forth motions of the tail (waggles) as the male attempted to lead females towards the pot. We defined pot entry as any time the focal male entered the half terra cotta pot at the center of his territory and digging as any time the male scooped gravel from inside its pot or around its pot into its mouth and subsequently released it around its pot. Stable DOM males exhibited changes in behavior from the pre-injection and BL period, but no changes between BL and the day of ascent (Supplementary Results and Figure S1). We took this as evidence of stable DOM males needing about a day to acclimate to the experimental tank to re-establish a territory in the new environment. Therefore, we only analyzed statistically behavior during the following periods: BL, morning of social ascent, and afternoon of social ascent. Videos were scored in Scorevideo (Matlab). The results of scoring videos were saved into log files that were subjected to a variety of analyses using custom R software (log files and code are available at: available at https://github.com/FernaldLab).

### Analyses of temporal patterns of behavior

### Interbehavior interval

The log files containing the results from Scorevideo were analyzed for interbehavior intervals (IBI)—the time between successive behaviors—using custom R software (available at https://github.com/FernaldLab). We determined average IBI during the morning and afternoon for DOM, ASC, and ASC+CA fish. ND fish performed almost no behavior so were not included in analyses of temporal patterns of behavior.

### Behavior sequence analysis

We conducted behavior sequence analysis using custom R software (available at https://github.com/FernaldLab). To determine the IBI needed to most accurately group successive behaviors into bouts for behavior sequence analysis, which we call “bout IBI”, we first determined the average IBI across DOM, ASC, and ASC+CA log files, regardless of treatment or time of day. This value (∼10 seconds) was used as bout IBI for behavior sequence analysis across DOM, ASC, and ASC+CA fish. Given ASC+CA fish had significantly longer IBIs compared to DOM and ASC fish (see Results), we conducted a second round of behavior sequence analysis for DOM, ASC, and ASC+CA fish using a bout IBI of 5.5 seconds, the average IBI across DOM and ASC files regardless of treatment or time of day. A third and fourth round of behavior sequence analysis was conducted on only DOM and ASC fish using bout IBIs of 5.5 seconds and 9 seconds, respectively (see Results). Behaviors that occurred outside of a given bout IBI were not included in behavior sequence analysis.

We quantified behavior sequences containing aggressive behavior (lateral displays and border fights) that were two behaviors in length in DOM, ASC, and ASC+CA males, since ASC+CA males basically performed no reproductive behavior (see Results for reasoning). For DOM and ASC males, we quantified behavior sequences containing aggressive and reproductive behavior that were two, three, four and five behaviors in length. Sequences containing six or more behaviors were too infrequent for meaningful analysis and were thus not analyzed.

For all behavior sequence analyses, we quantified the number of occurrences of a particular sequence and the transitional probabilities from specific behaviors. For instance, we counted the number of times the sequence “lateral display-chase female” occurred, but also quantified the probability of lateral display being followed by chase female. We also analyzed how behavior categories were sequenced using the same approach. For instance, we analyzed the transition probability of a male-directed aggressive behavior being followed by a reproductive or courtship behavior four behaviors later. When these analyses were conducted over sequences of four and five behaviors in length, we were only interested in the beginning and end of the behavior sequence of interest, so the types of behaviors in between were not considered and thus replaced with an “N” in the figures of results, where N=any behavior. Categories were defined as such: Male-directed aggressive behavior (lateral display and border fight), reproductive behavior (dig, pot entry, chase female, quiver, and lead swim), and courtship behavior (chase female, quiver, and lead swim). Sequence analysis was conducted using custom R software (available at https://github.com/FernaldLab).

For behavior sequences that were subjected to sequence analysis, sometimes a fish did not perform a component behavior. For instance, when analyzing the occurrence of the behavior sequence “lateral display-chase female” a fish may not have performed chase female. Because quantifying the occurrence of a behavior sequence in a fish that did not perform a component behavior may be confounded, fish that did not perform a component behavior of a behavior sequence were excluded from the analysis of that particular behavior sequence. Excluded fish for this reason occurred very infrequently and for any analysis when it did a maximum of two fish had to be excluded. The sample sizes for each test are shown in the Results section.

### Morphological and steroid hormone analyses

Focal fish were assessed for standard length (SL), body mass (BM), testes mass, and gonadosomatic index [GSI = (gonad mass/body mass)*100]. Blood samples were also collected. with capillary tubes from the caudal vein, centrifuged for 10 min at 5200 g, and the plasma was removed and stored at −80 °C until assayed. Immediately after blood collection fish were killed by cervical transection and brains were removed and immediately placed in a vial of PFA. Testes were removed and weighed to calculate GSI. For one ASC+CA fish, data for SL, BM, and testes mass could not be recorded (ASC+CA SL, BM, testes mass, and GSI final N=8).

Plasma testosterone and 11-ketotestosterone (11-KT) levels were measured using commercially available enzyme immunoassay (EIA) kits (Cayman Chemical Company, Ann Arbor, MI, USA) as previously described and validated for this species (Alcazar et al., 2016). Briefly, for testosterone and 11-KT assays, a 5-ul sample of plasma from each subject was extracted two times using 200 ul of ethyl ether and evaporated under a fume hood before re-constitution in EIA assay buffer (dilution 1:40 to 1:50). EIA kit protocols were then strictly followed, plates were read at 405 nm using a microplate reader (UVmax Microplate Reader; Molecular Devices, Sunnyvale, CA, USA) and steroid concentrations were determined based on standard curves. All samples were assayed in duplicate and intra-assay coefficients of variation (CV) were: testosterone (15.2%); 11-KT (8.9%). One DOM sample could not be assayed for testosterone due to possible contamination (DOM testosterone concentration final N=6), one ASC+CA plasma sample was not collected after the experiment, and 11-KT concentrations could not be determined for an ASC+CA fish due to limited sample (ASC+CA testosterone concentration final N=8; ASC+CA 11-KT concentration final N=7).

### Statistical analysis

All statistical tests were performed in the R statistical computing environment or Prism 7.0. We used Kruskal-Wallis ANOVAs followed by Dunn’s post-hoc tests for comparisons of physiological and behavioral measures across groups. To perform within-subject comparisons, we used Friedman’s Tests followed by Dunn’s post-hoc tests. When comparing only two groups, we used Mann-Whitney tests (between-subjects) or Wilcoxon tests (within-subjects). Raster plots were generated using custom software pacakges in R (available at https://github.com/FernaldLab). IBI analysis and behavior sequence analysis were also conducted using custom R software (available at https://github.com/FernaldLab). Correlational analyses were conducted using Spearmen’s rho. Effects were considered significant at p≤0.05.

## Results

### 1. ASC and ASC+CA males have similar circulating sex steroid hormones as stable DOM males

We found that the DOM, ASC, ASC+CA, and ND males used in this study were similar in measures of body size (Figure S2A; standard length, Kruskal-Wallis ANOVA, p=0.57; Figure S2B; body mass, Kruskal-Wallis ANOVA, p=0.74; N=6-8 per group). As expected, DOM males had significantly larger gonads (Figure S2C; Kruskal-Wallis ANOVA, p=0.0018; N=6-8 per group) and higher gonadosomatic indices [(gonad mass/body weight)x100] (Figure S2D; Kruskal-Wallis ANOVA, p=0.0015; N=6-8 per group) compared to the other groups. DOM, ASC, and ASC+CA males had higher levels of testosterone than ND males and did not differ from one another (Figure S2E; Kruskal-Wallis ANOVA, p=0.0041; N=6-8 per group). Circulating levels of 11-KT (Figure S2F; Kruskal-Wallis ANOVA, p=0.0006; N=6-7 per group) were significantly higher in DOM and ASC males compared to ND males (Figure S2F). DOM males also had higher levels of 11-KT compared to ASC+CA fish, which did not differ significantly from ASC or ND males. These results are in line with previous observations (Maruska et al., 2013; Maruska and Fernald, 2010; Parikh et al., 2006).

### 2. Androgen signaling is required for enhancing reproductive—but not aggressive— behavior during social ascent

We analyzed a panel of behaviors encompassing multiple categories (Fernald and Hirata, 1977; examples are in Figure 1B), including subordinate behavior, fleeing, where the male flees from a stimulus male; male-directed aggressive behavior, such as border fights and lateral displays; female-directed aggressive behavior, where the male attacks a stimulus female; and reproductive behavior, including chase female, quiver at female, lead swim, pot entry, and digging. These behaviors are described in detail in the Methods.

Behavior across groups on the day of ascent was visualized in raster plots to assess qualitative differences (Figure 3). Representative plots from each group show DOM, ASC, and ASC+CA males performed similarly high levels of aggressive behavior in the morning and afternoon, while ND performed almost no aggressive behavior. DOM and ASC males performed similarly high levels of reproductive behavior in the morning and afternoon, while ASC+CA and ND males performed virtually no reproductive behavior.

**Figure 3.**
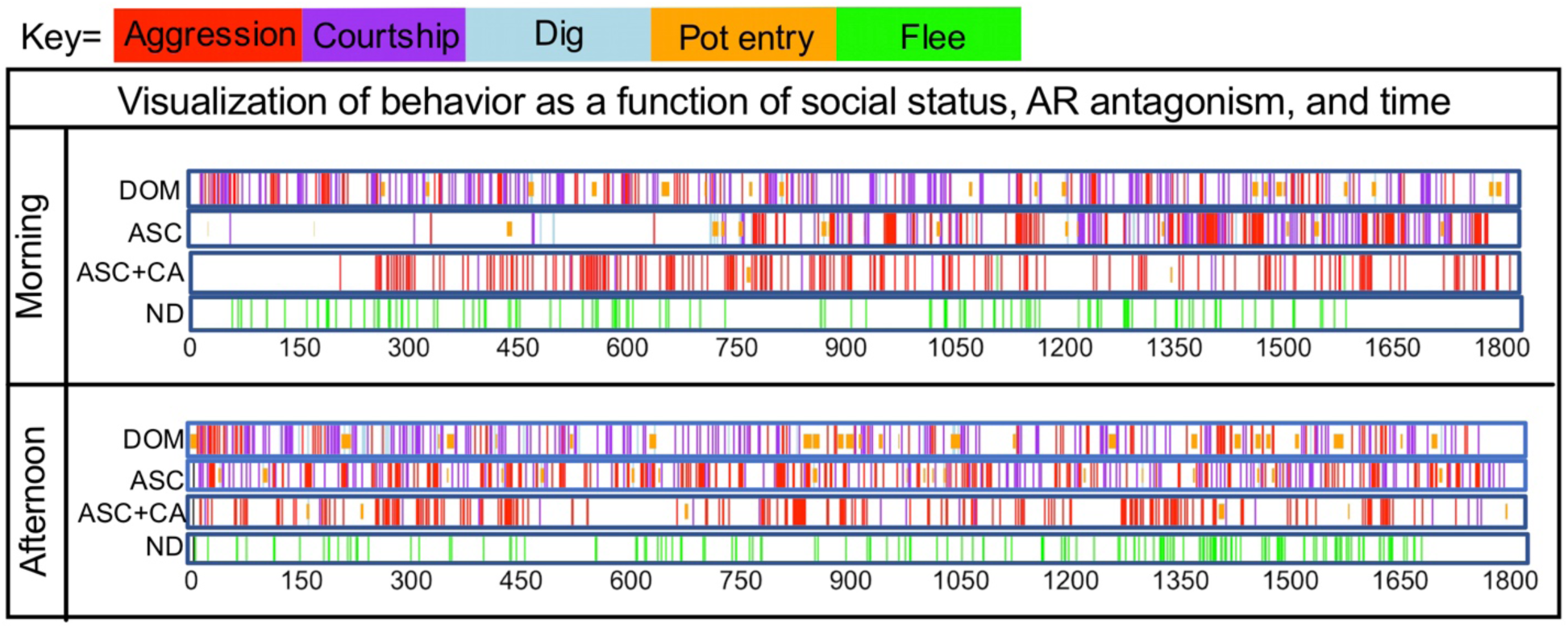
Raster plots showing behavior from individual fish from each group in the morning and afternoon. Each colored line represents a particular type of behavior. The x-axis represents time. Pot entry here is designated as durational; once the bar denoting pot entry is over the fish has exited the pot.

Given these observations, we next queried the extent to which there are quantitative differences between ASC and ASC+CA during ascent by analyzing changes in behavior from BL to the morning and afternoon on the day of ascent. ASC and ASC+CA males performed more aggressive behavior on the day of ascent compared to baseline (Figure 4A and Figure S3A; Friedman’s ANOVAs, p<0.05 for both groups, N=7-8 per group). Neither group showed a change in attacks directed towards females (Figure S3; p≥0.41 for both tests; N=7-8 per group). ASC males also significantly reduced fleeing behavior (Figure S3A; p=0.0014; N=7). For ASC+CA males, there was a non-significant decrease in fleeing behavior from baseline to day of ascent (Figure S3B; p=0.08; N=8). However, while ASC males showed a significant increase in reproductive behaviors compared to baseline (Figure 4A and Figure S3A; p<0.05, N=7), ASC+CA males did not (Figure 4B and Figure S3B; p≥0.12 for all tests; N=8), suggesting androgen signaling is required for a key aspect of social ascent, an increase in the performance of reproductive behavior.

**Figure 4.**
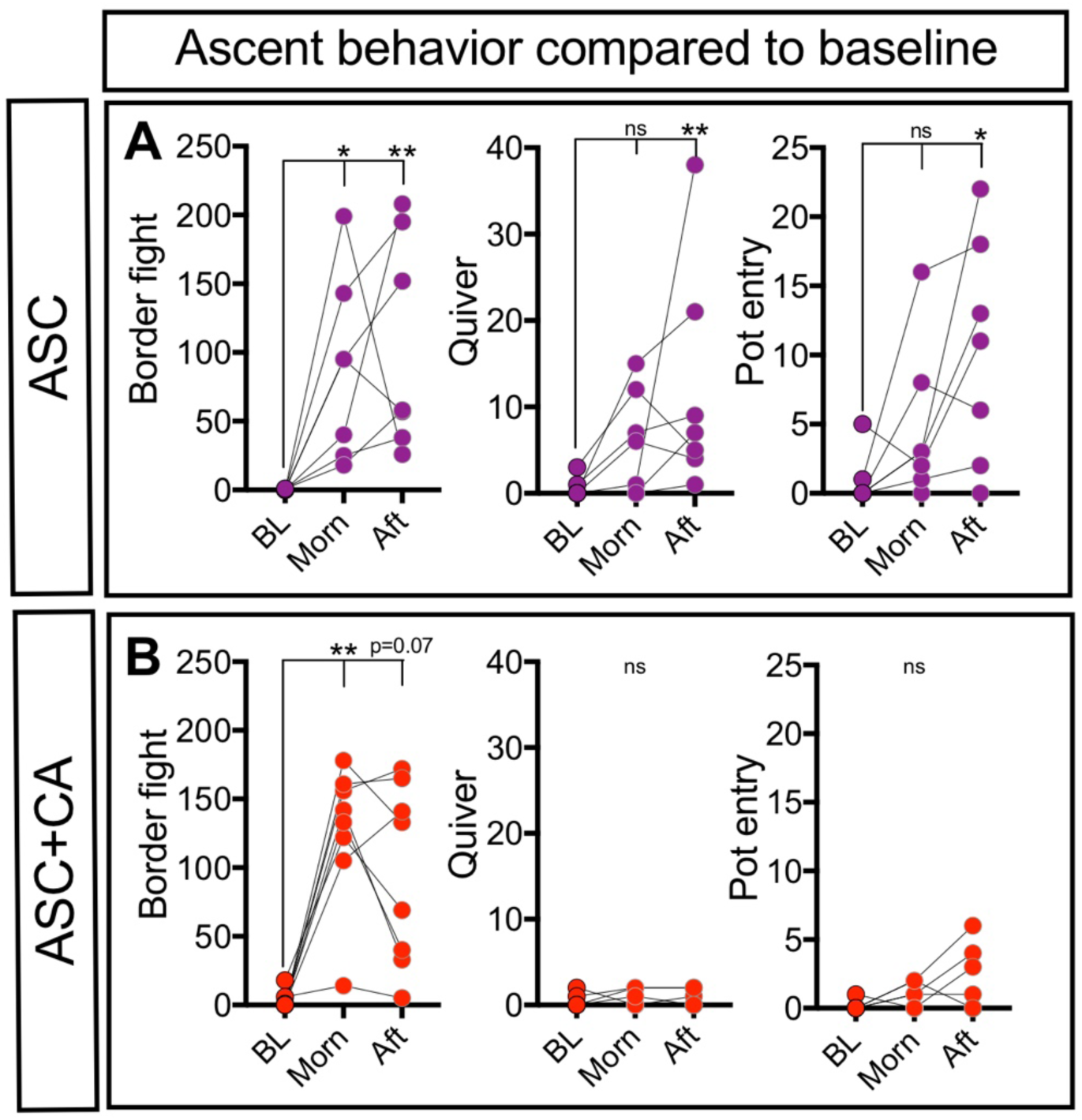
Plots showing change in a variety of behaviors from BL to the morning and afternoon on the day of social ascent for ASC and (B) ASC+CA fish. ns=non-significant; *p<0.05, **p<0.01. ASC=ascender; CA=cyproterone acetate; BL=baseline; morn=Morning; Aft=Afternoon.

Do ASC or ASC+CA male behaviors differ on the day of social ascent compared to stable DOM and ND males? In the morning ASC and ASC+CA males performed DOM-typical levels of aggressive behavior (Figure 5A and Figure S4A; Kruskal Wallis ANOVAs, p<0.05 for all main effects; N=6-8 per group). There was a main effect of social status on the number of attacks directed towards females (Figure S4A; Kruskal Wallis ANOVA, p=0.04; N=6-8 per group), where ASC+CA males attacked females more than ND males; however, no other differences between groups were observed. ASC males performed levels of reproductive behavior that were not different than levels seen in DOM males; however, they also did not differ statistically from ASC+CA or ND males. However, ASC+CA males performed virtually no reproductive behavior and did not differ from ND males (Figure 5A and Figure S4A), further supporting the notion that androgen signaling is required for the performance of DOM-typical levels of reproductive behavior.

**Figure 5.**
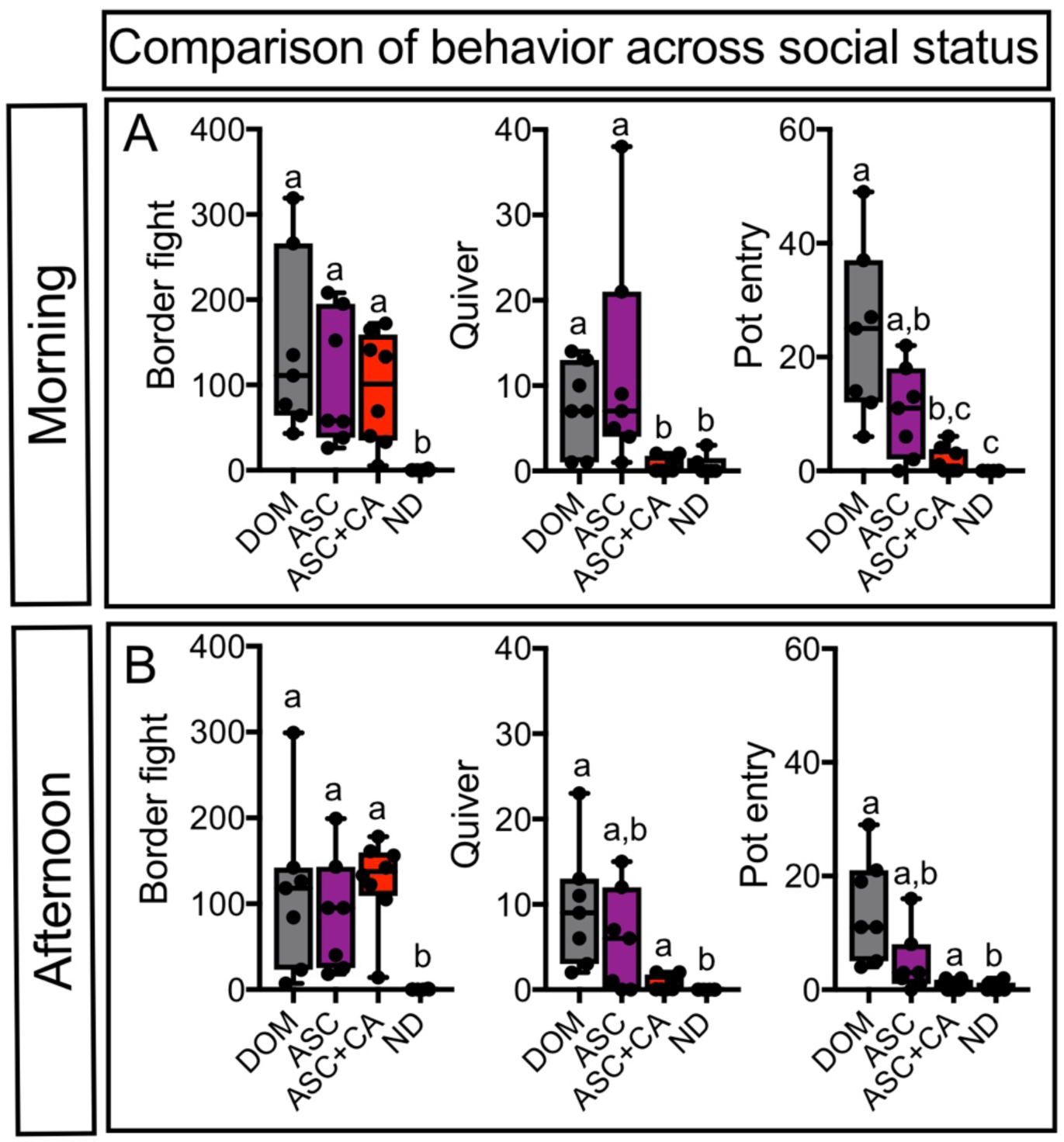
Box plots showing numbers of different behavior in the (A) morning and (B) afternoon performed by DOM, ASC, ASC+CA, and ND fish. Letters indicate if a group was statistically different (different letters) or statistically indistinguishable (same letters) from the other groups. DOM=dominant; ASC=ascending; ND=non-dominant; CA=cyproterone acetate.

Overall, in the afternoon the pattern of behavioral differences was similar to the morning (Figure 5B and Figure S4B, and Supplementary Results), but some differences emerged. For instance, while ASC and ASC+CA males differed statistically from ND males in the number of lateral displays performed, DOM males did not differ statistically from ASC, ASC+CA, or— strikingly—ND males (Kruskal Wallis ANOVA, p=0.0013; Figure S4B). This surprising lack of a difference between DOM males and the other groups in the afternoon prompted us to analyze temporal changes in the performance of lateral displays in DOM males. We found that DOM males performed significantly fewer lateral displays over time (Median difference between morning and afternoon: −18; Wilcoxon test, p=0.01; N=7; data not graphed). Overall then, our findings suggest that while androgen signaling in ASC males is required for the performance of DOM-typical levels of reproductive behavior, it is not sufficient for the expression of DOM-typical temporal behavioral patterns.

### 3. Social ascent is characterized by AR-dependent—and AR-independent—changes in temporal patterns of behavior

### 3.1 Time between successive behaviors is AR-dependent

After finding that differences between the groups changed over the course of the day, we next asked if blocking androgen signaling in socially ascending fish disrupts the speed at which behaviors are performed, which can be measured using inter-behavior interval (IBI). We measured the IBI in the morning and afternoon by measuring the interval between successive behaviors, and computed the average of this measure for DOM, ASC, and ASC+CA fish. ND males were not included in this analysis because they performed virtually no behavior. DOM, ASC, and ASC+CA fish had similar IBI in the morning (Figure S5A). In the afternoon, however, ASC+CA fish performed behavior at significantly slower speed (i.e., with significantly longer IBI) than DOM fish (Kruskal Wallis ANOVA, p=0.0097; Figure 6A). ASC fish were not different from either DOM or ASC+CA fish (p=0.19 and 0.91, respectively). These data imply androgen signaling in socially ascending fish is required for enhancing the speed of behavior performance, not just the occurrence of individual reproductive behaviors.

**Figure 6.**
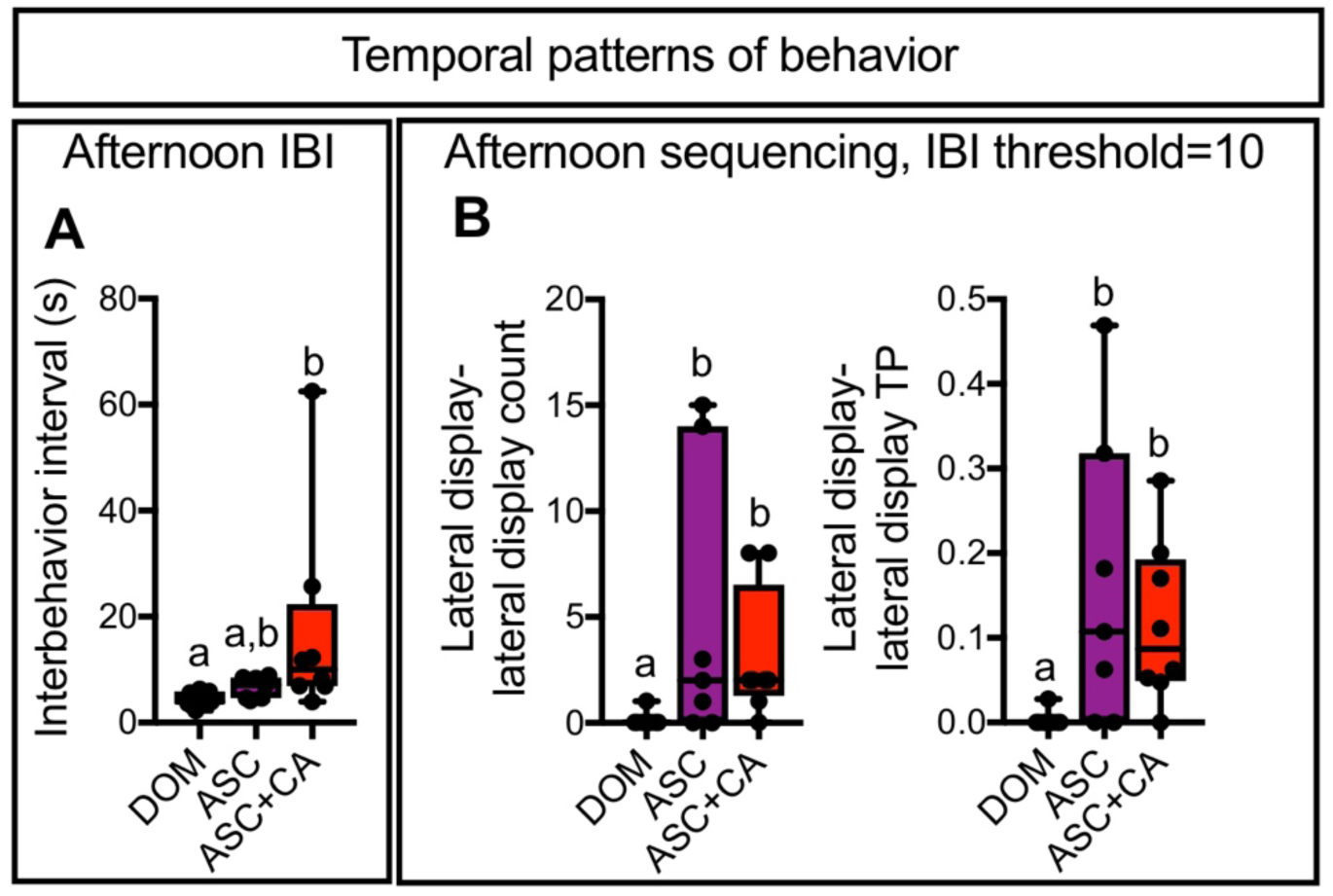
Box plots showing (A) IBI and (B) behavior sequencing across DOM, ASC, and ASC+CA fish. Letters indicate if a group was statistically different (different letters) or statistically indistinguishable (same letters) from the other groups. DOM=dominant; ASC=ascender; IBI=interbehavior interval; Count=number of times sequence occurred; TP=transition probability, the probability a given behavior followed an initial behavior.

#### 3.2.1 Lateral displays are repeated more often by ASC regardless of AR activation

We also wondered whether androgen signaling in socially ascending fish controls the sequencing of behavior. Since ASC+CA fish performed virtually no reproductive behavior, we only quantified the sequencing of aggression and compared it to DOM and ASC fish. We analyzed behavioral sequences within bouts of behavior, which were defined as two or more behaviors separated by 10 seconds or less, the average IBI across all DOM, ASC, and ASC+CA fish regardless of time of day (See Methods). There were no significant differences in the sequencing of aggression in the morning (Kruskal Wallis ANOVAs, N=7-8, p≥0.09 for all comparisons). However, in the afternoon we found a significant effect of social status—but not CA treatment— on the sequencing of male-directed aggressive behavior (Kruskal Wallis ANOVAs, N=7-8, p<0.05). Specifically, ASC and ASC+CA fish performed lateral displays twice in a row more often than DOM fish (Figure 6B; Dunn’s test, p<0.05 for both comparisons). ASC and ASC+CA males were also more likely to follow a lateral display with another lateral display compared to DOM males (Figure 6B; Dunn’s test, p<0.05 for both comparisons).

We conducted the above comparisons again, except we defined bouts of behavior as two or more behaviors separated by 5.5 seconds or less, the average IBI across DOM and ASC fish regardless of time of day (See Methods). This did not alter the general pattern of results (Figure S5B), indicating differences in behavioral sequencing are not due to differences in the tempo of behavioral performance. DOM, ASC, and ASC+CA fish performed similar numbers of lateral displays in the afternoon, suggesting differences in temporal patterning of lateral displays cannot simply be attributed to differences in the number of individual lateral displays performed (Figure S4). These results as a whole suggest another temporal dimension of behavior—how behaviors are sequenced—demarcates socially ascending fish from stable DOM fish.

#### 3.2.2 Ordering of successive behaviors differs between DOM and ASC in the afternoon

Given our observation that sequencing patterns of aggressive behavior demarcate socially ascending versus stable DOM male *A. burtoni*—regardless of AR activation—we wondered how the full suite of DOM-typical behaviors were sequenced differently in ASC and DOM males. ASC+CA males were excluded from this analysis because they performed virtually no reproductive behavior. We set the acceptable time interval between behaviors to be analyzed as 5.5 seconds and measured behavioral sequences comprising two, three, four, and five behaviors (See Methods).

DOM and ASC male behavioral sequences did not differ in the morning (Mann-Whitney test, p≥0.09 for all comparisons of behavior sequences; N=7 for both groups). However, there were striking differences in behavior sequences between DOM and ASC males in the afternoon on the day of ascent. DOM males were more likely to follow lateral displays with female chases and performed this sequence more often than ASC males (Figure 7A and Figure S5; Mann-Whitney test, p<0.05 for both comparisons; N=7 for both groups). DOM males were more likely to perform a lead swim after exiting their pot and performed this sequence more often than ASC males (Figure 7A and Figure S6; p<0.05 for both comparisons; N=5-7). We also asked how different categories (i.e., male-directed aggression, reproductive, and courtship behavior) of behavior were sequenced and found that ASC males were more likely to follow male-directed aggression with male-directed aggression than DOM males (Figure S6). Similarly, there was a statistical trend wherein DOM males were more likely to follow male-directed aggression with a reproductive behavior than ASC males (Figure S6; p=0.07 for both; N=7 for both groups). Thus, androgen signaling in ASC males is not sufficient for activating DOM-typical sequencing patterns of aggressive and reproductive behavior, suggesting another temporal aspect of behavior demarcates ASC and DOM social status.

**Figure 7.**
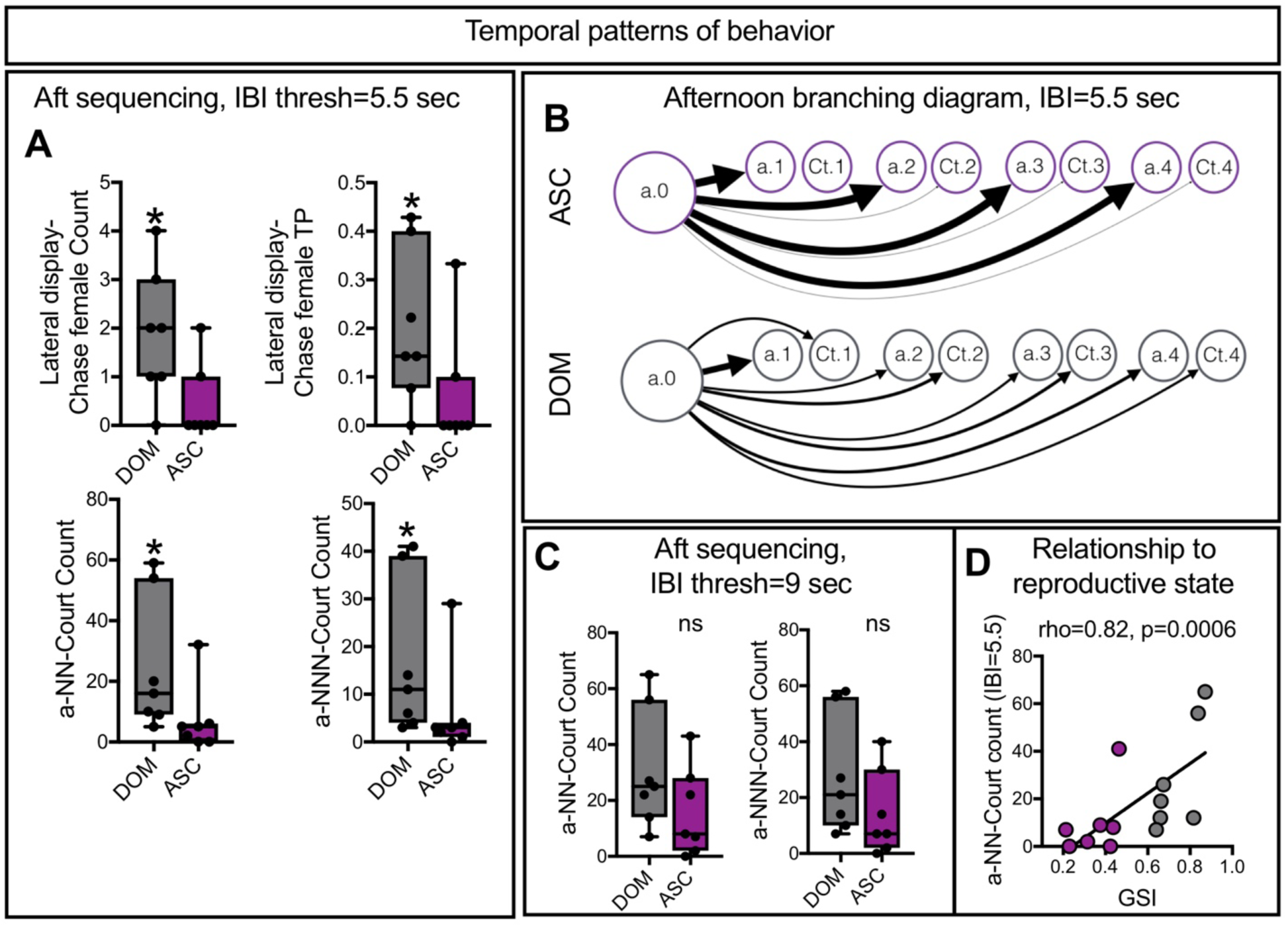
Behavioral sequencing patterns in the afternoon using a (A) 5.5 sec bout IBI and 9 sec bout IBI. (C) Relationship between gonadosomatic index (GSI) and a behavior sequence of four behaviors in length. (D) Behavioral sequencing patterns following male-directed aggression to male-directed aggression or courtship (i.e., a branching diagram) for a representative ASC and DOM fish. The largest arrow=53 occurrences; the smallest arrow=1 occurrence; for the ASC fish, there were no occurrences from a.0 to Ct.1 (hence, there is no arrow connecting a.0 to Ct.1. DOM=dominant; ASC=ascender; a=male-directed aggression; N=any behavior; IBI=inter-behavior interval; sec=seconds. Ct=Courtship behavior.; Count=number of times sequence occurred; TP=transition probability (i.e., the probability a behavior followed an initial behavior). *p<0.05; ns=non-significant.

### 3.3 Longer behavioral sequences distinguish DOM from ASC males

Beyond the ordering of two successive behaviors, we next asked whether longer behavioral sequences differed between groups (i.e. sequences > 2 behaviors) and found that, for multiple sequence lengths, many differences associated with male-directed aggressive behaviors emerged in the afternoon. In three-behavior sequences, the same general pattern as for two-behavior sequences remained, but some new differences emerged (Figure 7 and Figure S6). DOM males, after performing two border fights successively, were more likely to perform a female chase compared to ASC males (Mann-Whitney test, p=0.03; N=7 for both groups). This pattern persisted when behavioral categories were analyzed (Figure 7A and Figure S6). For instance, after performing two male-directed aggressive behaviors, DOM males were more likely to perform a reproductive or courtship behavior compared to ASC males (Mann-Whitney test, p<0.05 for both comparisons; N=7 for both groups). Strikingly, this pattern of differences persisted for sequences that included four and five behaviors: DOM males performed more behavior sequences that started with male-directed aggression and culminated in reproductive or courtship behavior three and four behaviors later (Figure 7A-B and Figure S6; Mann-Whitney test, p<0.05 for both comparisons; N=7 for both groups).

Thus, ASC- and DOM-typical behavior sequencing patterns are detectable in short and long behavior sequences. Specifically, ASC males typically follow aggressive behavior with more aggressive behavior, while DOM males typically follow aggressive behavior with reproductive or courtship behavior. Therefore, androgen signaling in ASC males is not sufficient for activating DOM-typical sequencing patterns of aggressive and reproductive behavior, suggesting yet another complex distinction between ASC and DOM social status. Our findings on behavioral sequences could not be attributed to overall behavioral frequency, which we addressed in several ways (see Supplemental Results and Figure S7-S10).

### 3.4. A longer IBI eliminates certain transition differences

The differences in behavioral sequencing between DOM and ASC males might depend, in part, on the time used for defining a bout. For instance, it could be that ASC and DOM males perform courtship behavior after male-directed aggression at similar rates, but ASC males take longer than DOM males to transition from male-directed aggression to courtship. Half of the ASC males had an IBI approximately 2 seconds or more longer than the bout IBI used of 5.5 seconds and the highest IBI in the ASC group was ∼9 seconds. Thus, we conducted the behavioral sequence analysis again but used an IBI of 9 seconds instead of 5.5. Using a 9-second instead of a 5.5-second interval did not alter the general pattern of differences observed between the groups when behavior chains that were two and three behaviors long were considered (Table S2). However, differences between DOM and ASC were no longer present for behavior sequences that were four and five behaviors long (Figure 7C and Table S2). Therefore, the differences in behavioral sequences between DOM and ASC males are in general stable for short sequences, but differences in longer sequences as a function of status are eliminated when longer functional bout intervals are considered. These results suggest ASC males can perform some DOM-typical behavior sequences, but may take more time between successive behaviors.

### 4. Behavioral sequencing correlates with reproductive state

Finally, we tested whether the differences in behavioral sequencing in ASC and DOM males in the afternoon were related to reproductive state. Hence, we correlated the significantly different behavioral sequences to GSI and testes mass. Numerous significant linear correlations were observed (Table 1). The strongest positive correlation was between GSI and the occurrence of a chain of four behaviors, in which male-directed aggression was followed by courtship behavior three behaviors later (rho=0.79, p=0.0012; Figure 7D). These data indicate reproductive state is linearly related to the sequencing of behavior.

**Table 1.** Correlations between different behavior sequences and GSI and testes mass. d=lateral display; C=chase female; a=male-directed aggression; Repro=reproductive behavior; Court=courtship behavior; N=any behavior; x=pot exit; s=lead swim; b=border fight; Count=number of times the chain actually occurred; TP=transition probability. *significant correlation at denoted p value.

| **Table 1-Correlations to gonadal measures. N=14.** |  |  |  |  |
| --- | --- | --- | --- | --- |
|  | **GSI** |  | **Testes Mass** |  |
| **Behavior** | **rho** | **P** | **rho** | **P** |
| **dd Count** | -0.53 | 0.054 | -0.41 | 0.14 |
| **dd TP** | -0.6 | 0.025* | -0.43 | 0.12 |
| **dC Count** | 0.76 | 0.0025** | 0.56 | 0.04* |
| **dC TP** | 0.63 | 0.01* | 0.48 | 0.08 |
| **xs Count** | 0.65 | 0.014* | 0.54 | 0.04 |
| **xs TP** | 0.62 | 0.02* | 0.55 | 0.04* |
| **aaRepro TP** | 0.51 | 0.06 | 0.63 | 0.02* |
| **aaCourt TP** | 0.6 | 0.02* | 0.65 | 0.01* |
| **bbC TP** | 0.67 | 0.01* | 0.56 | 0.04 |
| **a-NN-Repro Count** | 0.68 | 0.0093** | 0.49 | 0.07 |
| **a-NN-Court Count** | 0.79 | 0.0012** | 0.49 | 0.07 |
| **a-NNN-Repro Count** | 0.69 | 0.0084** | 0.49 | 0.07 |
| **a-NNN-Court Count** | 0.75 | 0.003** | 0.53 | 0.05* |

## Discussion

We used a social ascent paradigm in *A. burtoni* combined with AR antagonism to dissect the suite of behavioral traits that characterize social ascent and dominance. We demonstrate that androgen signaling is required for the performance of reproductive behavior during social ascent. Moreover, we showed androgen signaling as well as social status influences the temporal patterning of behavior. Our pharmacological dissection of social ascent revealed androgen signaling and experience may interact to regulate social behavior in *A. burtoni,* and social status may be dissociable.

### Androgen signaling governs multiple aspects of the social calculus used by male *A. burtoni*

The roles played by sex steroid hormone signaling in regulating aggressive and reproductive behavior is widely appreciated (Eisenegger et al., 2011) and our findings are consistent with other studies on the regulation of courtship behavior by androgen signaling in teleost fish (Almeida et al., 2014; O’Connell and Hofmann, 2012a). However, the hormonal mechanisms underlying the likelihood of an animal rising in social rank are less clear.

Using *A. burtoni*, we showed that blocking androgen signaling virtually abolished all forms of reproductive behavior in males that were given the opportunity to rise in social rank. This differs markedly from previous work (O’Connell and Hofmann, 2012a) showing that in *A. burtoni*, blocking ARs in DOM males reduced only one aspect of courtship—quivers—and no other reproductive or aggressive behavior. Interestingly, ND males given exogenous androgens do not increase reproductive behavior (O’Connell and Hofmann, 2012a). Thus, our results suggest that AR has distinct regulatory effects during the rapid behavioral shift that occurs after perceiving social opportunity than it does during the behavior of stable DOM and ND males.

Indeed, it is well known that male *A. burtoni* survey the environment to guide their behavior (Desjardins et al., 2012; Grosenick et al., 2007), a process that has been called “social calculus” (Fernald, 2017). Our results suggest that this social calculus may require androgen signaling. This is in line with studies in humans (Dreher et al., 2016; Eisenegger et al., 2011), non-human primates (Lacreuse et al., 2010), rodents (Aikey et al., 2002; Frye et al., 2008; Jacome et al., 2016), and birds (Alward et al., 2016, 2017; Bottjer and Hewer, 1992) that show androgens affect complex cognitive processes (McCall and Singer, 2012). For instance, testosterone shifts player strategy to enhance social status in an experimental gambling paradigm (Dreher et al., 2016). Hence, AR antagonism may have led to deficits in the recognition of social opportunity (Berghe, 2014; Fernald, 2017). Multiple sensory modalities are likely involved in an ND male’s ability to recognize and act on social opportunity (Maruska and Fernald, 2012). AR is expressed in the retina of zebrafish (*Danio rerio*) (Gorelick et al., 2008) and brain regions that project to the olfactory bulb of *A. burtoni* (Harbott et al., 2007), suggesting androgens act in these areas to integrate a variety of chemosensory cues. ARs are also expressed in the POA of *A. burtoni*, a well-conserved brain region that integrates social cues and controls sexual behaviors across vertebrates (Goodson, 2005; Harbott et al., 2007; O’Connell and Hofmann, 2012b).

### AR-dependent and AR-independent temporal patterns of behavior delineate ASC from DOM males

While we expected AR blockade to reduce courtship behavior during social ascent, we observed surprising effects of AR blockade on social status and temporal patterns of behavior. Socially ascending males with blocked AR performed behavior at a slower speed (i.e., took more time between each successive behavior) than stable DOM males. Previous research has shown testosterone affects the timing of behavior in response to different cues. For instance, in rhesus monkeys (*Macaca mulatta*) testosterone treatment sped up reaction times to social cues during a task testing recognition memory, which may have reflected an enhancing effect of testosterone on reward sensitivity or general arousal (King et al., 2012). Testosterone treatment in rhesus monkeys also increased attention to videos depicting fights between conspecifics (Lacreuse et al., 2010). Given the well-established role of testosterone in mediating arousal and sexual motivation (Adkins-Regan, 2009; Eisenegger et al., 2011), it is intriguing to consider the possibility that blockade of androgen signaling in socially ascending male *A. burtoni* reduced reaction times to social cues, reflecting a state of reduced arousal and social motivation. These findings also mirror those found in songbirds, showing that AR blockade significantly slows down the speed at which sexually-relevant song is performed (Alward et al., 2016).

However, socially ascending fish—regardless of AR activation—sequenced aggressive behavior differently than stable DOM males. We further showed that DOM males are more likely than ASC males to follow aggressive behavior with courtship behavior, a difference that persisted for behavioral sequences of successively longer lengths. Some of these differences disappeared when we used a longer bout IBI, implying ASC males can perform some of the DOM-typical behavioral sequences, but they take longer to switch from male-directed aggression to female-directed courtship. This pattern of differences reflects an important dimension of social status, that to the best of our knowledge hitherto was not known.

### Differences in behavioral transitions may reflect AR- and experience-dependent social task switching: A working model

The rise in aggressive and reproductive behavior to DOM-like levels in socially ascending fish is likely mediated by neurons activated by chemosensory cues from neighboring males and females, respectively. One explanation for the disconnect between the performance of *individual* aggressive and reproductive behavior, the *speed* of behavior, and the *transitions* from aggression to aggression, or aggression to courtship, may be differences in neural responsiveness or selectivity for chemosensory cues from males and females in socially ascending males versus DOM males. This could manifest as a difference in how fast males behave in general and in how fast they transition from male-directed aggression to female-directed courtship behavior, if the decision or speed to transition is coordinated differently based on sensitivity to cues and/or social status. In other words, socially ascending and DOM males may differ in “social task switching”.

Sex steroid hormones and experience may interact to control an animal’s responsiveness to different cues. Indeed, multiple observations in different taxa support the idea that physiological or maturational state influences cognitive processes like task switching. In humans, performance on tests measuring task switching improves from childhood to puberty and adulthood, suggesting sex steroid hormones and/or experience are involved (Davidson et al., 2006). Pre-pubertal rats perform more perseverative behavior compared to post-pubertal and adult rats during the water maze task (Juraska and Willing, 2017). Critically, gonadectomized male and female rats performed poorly at a go/no-go task measuring urinary odor sex discrimination, but after receiving testosterone implants, both sexes performed significantly better (Kunkhyen et al., 2018).

Social interactions in *A. burtoni* may lead to long-term changes in steroid actions in key brain regions that shape neural selectivity to distinct social cues. Indeed, neuronal responses are tuned by experience or hormones. For instance, juvenile zebra finches during early stages of song learning possess neurons that respond to their own specie’s song (Solis and Doupe, 1997). However, these neurons do not discriminate between their own song and their tutor’s song until song is crystallized (Solis and Doupe, 1997), when birds experience an increase in circulating testosterone (Marler et al., 1988, 1987), the action of which has been shown to regulate aspects of song crystallization, including the temporal structure of song (Alward et al., 2017; Bottjer and Hewer, 1992; Korsia and Bottjer, 1991). Additionally, estradiol concentrations increase in the auditory cortex of male zebra finches when interacting with females or hearing another male’s song (Remage-Healey et al., 2008). The estradiol produced in the auditory cortex of zebra finches due to these interactions may underlie the enhancement in the selectivity to the bird’s own song of neurons in a downstream sensorimotor nucleus that controls song production (Remage-Healey and Joshi, 2012).

Taken together, it is attractive to consider the possibility that socially ascending males undergo piecemeal changes in neural responsiveness to male and female cues controlled by gonadal steroids and experience that eventually lead to the production of the full suite of DOM-like behavioral temporal patterns. Hence, in our working model (Figure 8), male-directed aggression requires the activation of ER+ neurons stimulated by chemosensory cues from neighboring males, a prediction based on studies by O’Connell and colleagues showing estrogen signaling activates aggression in *A. burtoni* (Huffman et al., 2013; O’Connell and Hofmann, 2012a). Our model also posits the performance of courtship behavior depends on the activation of AR+ neurons by female chemosensory cues (current findings and O’Connell and Hofmann, 2012a). The extent of this effect clearly depends on social status, as blockade of AR in stable DOM males in previous work only reduced quiver (O’Connell and Hofmann, 2012a), but we show blockade of AR in socially ascending fish abolishes all forms of reproductive behavior. Also included in our model is that activation of AR+ neurons leads to a faster tempo of behavioral performance, reflecting general arousal or motivation.

**Figure 8.**
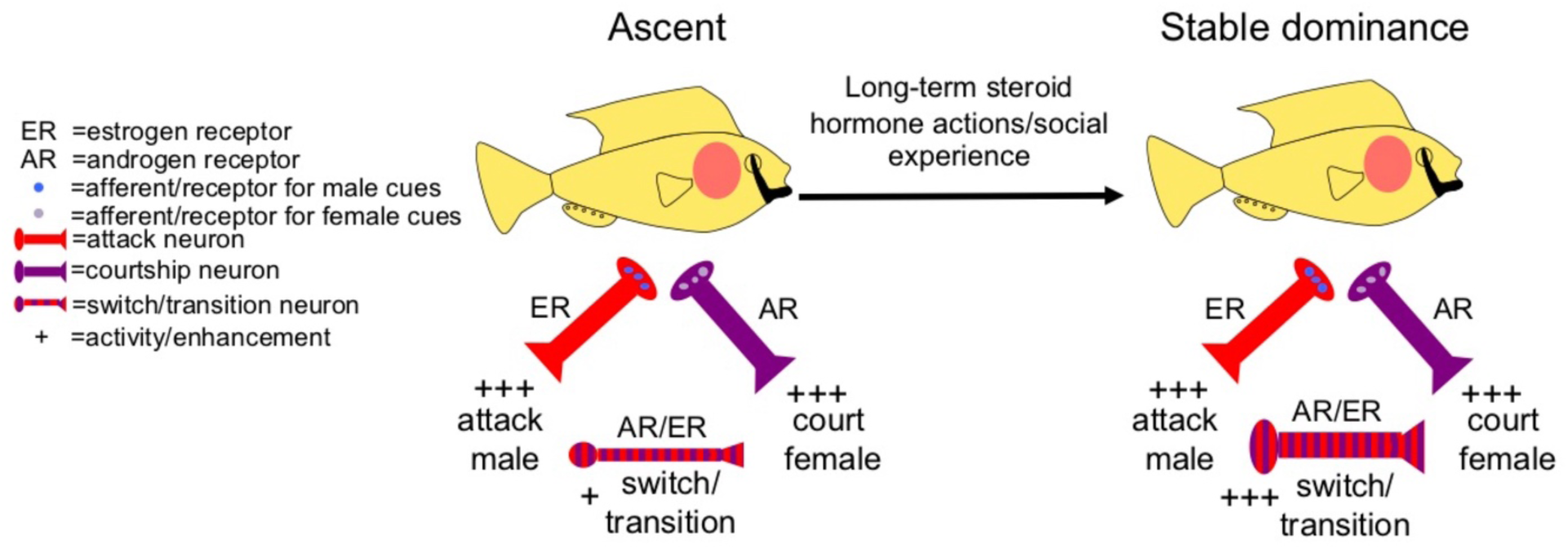
Working model on the regulation of social behavior in *A. burtoni*. When given the opportunity, ASC males increase aggression directed towards males through the activation of estrogen receptor (ER) positive neurons through afferent input about chemosensory cues from neighboring males. ASC males will also increase courtship directed towards females as a result of activation of androgen receptor (AR) positive neurons through afferent input about chemosensory cues from females. However, the neurons that modulate how males switch or transition from male-directed aggression to female-directed courtship are not as active or sensitive compared to DOM males, so these transitions occur less frequently (as is the case for lateral display-chase female) and take more time to take place (as is the case for long behavior chains that start with male-directed aggression and end with a reproductive or courtship behavior). These “switch” neurons may undergo a transformation in function after an ASC male is exposed to higher levels of sex steroid hormones and social experience for several days, wherein now males more rapidly transition from male-directed aggression to female-directed reproductive or courtship behavior. As discussed in the main text, one status is not merely more efficient at these transitions; it could be that for ASC males the decision to transition from male-directed aggression to female-directed courtship is weighted by factors such as the state of territory establishment and fight outcome more strongly than DOM males, which have established their territory. While this simplified schematic denotes each behavior as being controlled by specific neurons, it could be networks of neurons control these distinct behaviors. Our model presents tractable hypotheses that once tested may reveal fundamental aspects of the regulation of social behavior across species. Key: neuron size positively corresponds to hypothesized activity/sensitivity; the striped switch/transition neurons indicate it could integrate cues about attacking male and when to switch to courting females.

However, socially ascending males, regardless of AR blockade, are more likely than stable DOM males to follow male-directed aggression with male-directed aggression. Stable DOM males, on the other hand follow male-directed aggression more often with courtship behavior. These differences in behavioral sequencing patterns could be mediated by differential actions of gonadal steroids acting over different time scales. For instance, activation of AR+ neurons before/during social ascent is required to enhance the tempo of behavior overall to DOM-typical levels. However, given our results that ASC males—regardless of AR activation—do not perform the same behavioral sequence patterns as stable DOM males, it could be that the long-term exposure to sex steroid hormones is required in ASC males to enhance the rapid transitions from male-directed aggression to courtship in a DOM-typical manner. Transitions from male-directed aggression to courtship may be controlled by AR/ER+ neurons or neural networks that we will call “switch neurons”. In ASC males, these switch neurons do not discriminate as well or perform different computations between male and female chemosensory cues, leading to fewer or slower transitions from male-directed aggression to courtship. The switch from male-directed aggression to courtship may be controlled by interactions between attack-male neurons and courtship neurons or neurons that specifically control behavior switching (Figure 8). Our working model presents a foundation of testable hypotheses that are applicable to a wide-range of species and social behaviors, especially those that rely on experience.

## Conclusion: Pharmacological and behavioral dissection of social behavior reveals social status is highly dissociable

Steroid hormones are known to regulate physiology, morphology, and behavior at different time scales to coordinate an animal’s adaptive response to the environment (Adkins-Regan, 2009; Alward et al., 2017; Lee and Pfaff, 2008; Pfaff et al., 2008). Our results show that androgens may affect the complex social calculus used by animals as they determine their rank along a social hierarchy. Moreover, we provide evidence that gonadal steroids and social experience may also be involved in coordinating the temporal patterning of behavior, perhaps over distinct time scales or in integrating current behavior with experience. Our pharmacological and behavioral analyses revealed that the suite of behaviors that characterize social status are highly dissociable, suggesting non-redundant neural and hormonal mechanisms underlie an animal’s social decision making, which is reflected in the performance of individual behaviors and their temporal patterning. This type of control by gonadal steroids may be important in regulating complex social behaviors in other vertebrates.

## Acknowledgements

We thank Danielle Blakkan for help with the experimental set-up and injections and Paul Tran for scoring videos of the behavior.

**Author’s contributions:** Conceptualization, B.A.A. and R.D.F.; Methodology, B.A.A and R.D.F.; Investigation, B.A.A., A.T.H., and R.A.Y; Writing – Original Draft, B.A.A.; Writing – Review & Editing, B.A.A., R.D.F, A.T.H, and R.A.Y; Supervision, B.A.A. and R.D.F.

